# Prevalence and phenology of mycorrhizal colonization across populations of *Lycopodiella inundata*

**DOI:** 10.1101/2020.05.07.080192

**Authors:** Jill Kowal, Elena Arrigoni, Jordi Serra, Martin Bidartondo

**Affiliations:** Comparative Plant and Fungal Biology, Jodrell Laboratory, Royal Botanic Gardens, Kew, London TW9 3AB, UK; Department of Neuroscience, King’s College London, UK; Department of Life Sciences, Imperial College London, SW7 2AZ, UK

**Author notes:** Corresponding author: Jill Kowal.

**Keywords:** fine root endophytes (FRE), heathland ecology, Mucoromycotina, mycorrhizal phenology, plant-fungus interaction

## Abstract

Mycorrhizal fungi are critical components of terrestrial habitats and agroecosystems. Recently, Mucoromycotina fine root endophyte (MucFRE) fungi were found to engage in nutritional mutualism with the rare plant *Lycopodiella inundata* (‘marsh clubmoss’), one of the earliest vascular plant lineages known to associate with MucFRE. The extent to which this mutualism plays a role in resilient plant populations can only be understood by examining its occurrence rate and phenological patterns.

To test for prevalence and seasonality in colonization, we examined 1,297 individual *L. inundata* roots collected during spring and autumn 2019 from 11 semi-natural heathlands in Britain and the Netherlands. We quantified presence/absence of MucFRE-like hyphae and vesicles and explored possible relationships between temperature and precipitation in the months immediately before sampling.

MucFRE-like hyphae were the dominant mycorrhizal fungi observed in all of the examined heathlands. However, we found significant differences in colonization between the two seasons at every site. Overall, 14% of *L. inundata* roots were colonised in spring (2.4% with vesicles) compared with 86% in autumn (7.6% with vesicles). Colonization levels between populations were also significantly different, and correlated with temperature and precipitation, suggesting some local environments may be more conducive to hyphal growth.

These marked seasonal differences in host-plant colonization suggest that results about mycorrhizal status - typically drawn from single time point collections - should be carefully interpreted. Our findings are directly relevant to habitat restoration, species conservation plans, agricultural bio-inoculation nutrient enhancement treatments, microbial diversity and functional studies of host plants and symbionts.

## Introduction

Nutritional mutualistic exchange between mycorrhizal fungi and plant roots coevolved over millions of years (Pirozynski and Malloch 1975; Brundrett 2002; Bidartondo et al. 2011; Strullu-Derrien *et al*. 2014; Field et al. 2015a). This interaction is fundamental to plant resilience, particularly in stressful environments (Smith and Read 2010, Kowal et al. 2016), and ecosystem function above and below ground (Hart and Klironomos 2003; Giovannetti 2008). It is well established that endophytic fungi associate with early-diverging vascular plant genera such as *Lycopodium* (Winter and Friedman 2008; Imhof 2013) and *Lycopodiella* (Rimington et al. 2014). Importantly, *Lycopodiella* was recently found to engage in a nutritional mutualism with Mucoromycotina FRE (MucFRE) (Hoysted et al. 2019). MucFRE are the only mycorrhizal fungi to have been detected *in L. inundata* roots (Rimington et al. 2014; Hoysted et al. 2019), in sharp contrast with other vascular plants where fine root endophytes (FRE) and Glomeromycota arbuscular mycorrhizal fungi (AMF) coexist. This provides a unique opportunity to study exclusive FRE colonization in a vascular plant.

*Lycopodiella inundata* (L.) Holub (marsh clubmoss) is a rare herbaceous lycopod (Garcia Criado et al. 2017) found in nutrient-poor wet habitats across the Northern Hemisphere (Hultan and Fries 1986). In Britain and Europe, *L. inundata* associates with seasonally inundated heathland vegetation (Byfield and Stewart 2008), often along tracks and at the edges of oligotrophic lakes (Smyth et al. 2015; Korzeniak and Onete 2016; Price 2019). The plant’s spore-driven life history alternates between two generations - gametophytic and sporophytic. The sporophyte’s mature strobili produce spores in the late summer, resulting in diminutive gametophytes from late summer to spring. The rhizomatous stems spread mainly by creeping axes which can also successfully reproduce through vegetative fragmentation (Byfield and Stewart 2008). Dry hot summers and freezing winter temperatures influence both the degree to which the stem will die back above ground and/or continue to produce new roots below ground. However, the extent and necessity for fungal symbiosis in mature wild populations of *L. inundata* is unknown.

Fine root endophytes have a global distribution and are important in both agricultural and semi-natural systems across a broad range of host plant families (Field et al. 2015a, Orchard et al. 2017a). FRE were once considered a single species - *Glomus tenue*, basionym *Rhizophagus tenuis* (Orchard et al. 2017b) - but are now recognized as a group of taxa in the subphylum Mucoromycotina (Orchard et al. 2017a) within the genus *Planticonsortium* C. Walker et D. Redecker gen. nov. (Walker et al. 2018).

Here we investigated the prevalence of FRE colonization by examining large, geographically diverse populations of *L. inundata* sporophyte roots to determine whether they are ubiquitously and/or preferentially associating with FRE. There have been only a few reports specifically studying FRE colonization during a plant’s growth season (Thippayarugs et al. 1999; Fuchs and Haselwandter 2004; Bueno de Mesquita et al. 2018). Thus, by examining roots in the beginning of the growth season (spring) and six months later at the end (autumn) we aim to address the question of seasonality. We hypothesized that there are distinct seasonal patterns in FRE on a par with phenological developments in the host plant as well as differences between sites based on local climate variations. Moreover, by studying FRE phenology in the roots of this ancient plant lineage, we can gain further insight into abiotic and environmental drivers of this important fungal group. We also aim to begin isolating factors affecting retention of contracting *L. inundata* populations, informing conservation and restoration plans.

## Methods

### Site selection and sampling

Based on the population distribution of *L. inundata*, we selected six lowland UK heathlands and one metapopulation in the Netherlands, providing a west to east oceanic climate gradient. We also selected three northern Scotland heathlands to maximally contrast precipitation and temperature. Site visits were ordered according to latitude and scheduled as close together as possible (Table 1). Spring collections occurred over eight weeks commencing mid-April 2019 and autumn collections over six weeks commencing mid-September 2019.

**Table 1.** Sites and associated dates of *Lycopodiella inundata* sample collections.

| Site | Abbr. name | Owner/ Manager | Conservation status | Grid reference | County Country | Spring date | Autumn date |
| --- | --- | --- | --- | --- | --- | --- | --- |
| Hyde Bog | HB | Forestry Commission | SSSI National Nature Reserve | SY 87XXX 91996 | Dorset England | 15/04/19 | 16/09/19 |
| Park Lake | PL | Private | Not evaluated | SX XXXXX XXXXX | Cornwall England | 16/04/19 | 09/09/19 |
| Stannon Park | SP | Private | Not evaluated | SX XXXXX XXXXX | Cornwall England | 16/04/19 | 09/09/19 |
| Stadbury Hill | SH | National Trust | New Forest National Park | SU 28XXX 15959 | Hampshire England | 24/04/19 | 02/10/19 |
| Matley Heath | MH | Forestry Commission | New Forest National Park | SU 33XXX 08479 | Hampshire England | 24/04/19 | 02/10/19 |
| Thursley Common | TH | Natural England | SSSI National Nature Reserve | SU 90XXX 40827 | Surrey England | 04/05/19 | 18/09/19 |
| Aldershot | AL | MOD/Hampshire & Isle of Wight Wildlife Trust | Special Protection Area, EU | SU 83XXX 52511 | Hampshire England | 04/05/19 | 18/09/19 |
| Strabrechtse Heide | NL | Brabants Landschap/ Staatsbosbeheer | 2006 Birds Directive, EU; Natura 2000 | N 51 23' 48.16212'<br>E 5 35' 39.53544' | Heeze Netherlands | 27/05/19 | 11/10/19 |
| Munday Estate | MP | Private | Not evaluated | NG 81XXX 52419 | Wester Ross Scotland | 11/06/19 | 29/10/19 |
| Coulin Estate | CE | Private | Not evaluated | NG 98XXX 57635 | Wester Ross Scotland | 11/06/19 | 29/10/19 |
| Beinn Damh | BD | Private | Not evaluated | NG 85XXX 54609 | Wester Ross Scotland | 12/06/19 | 30/10/19 |

We generated three to five random 1m^2^ subplots depending on area covered by the *L. inundata* population. At Munday Path (MP), Beinn Damh Estate (BD) and Aldershot (AL), where the area of the site was too small to differentiate 1m^2^ subplots, we sampled from three plant clusters as far apart as possible from each other. We collected six *L. inundata* plants from each 1 m^2^ subplot (or cluster at MP and BD) but only three from AL, where the population size has rapidly declined in recent years. Care was taken to collect only from plants with photosynthetic tissue appearing healthy without decay.

Collections were repeated spring and autumn of 2019 within the sample population footprint and time schedule, but new subplots (or clusters) were used to avoid over-collecting.

### Root processing and analysis

Field collected plants were placed in cold storage (4°C) with their soil intact. To minimize under-detection of FRE, samples were processed within three days of collection (Orchard et al. 2017a,c). Soil was loosened from the plant roots by intermittently soaking and placing them under running tap water taking care the roots stayed intact. Remaining soil was gently brushed away from the roots with a soft paintbrush.

#### Measurements of root per plant, root density and root length

We measured plant length, root length and root density (number of roots per plant cm) (autumn only) of the intact specimen (Supplemental Table 1).

#### Root staining protocol

After measuring, roots were cross-sectioned along the elongation zone and placed in a 2ml microcentrifuge tube (three individual root sections per tube) containing 70% ethanol for staining and microscopic examination.

We modified existing staining protocols (Vierheilig et al. 1998; Wilkes et al. 2019) as follows. The roots were cleared by boiling them in 10% KOH for 20 minutes and a further 30 minutes at 60°C. After rinsing 3x in dH_2_0, roots were stained in a 10% Shaeffer ink + 25% glacial acetic acid solution at 100°C for three minutes. Without further rinsing, roots were left overnight to de-stain in 1% acetic acid. We prepared semi-permanent slides (76 × 26mm / 0.8 - 1.0mm) by placing 200µl of 50% glycerol solution on the slide, adding 1-2 roots in the droplet, placing a cover slip (18mm^2^ / 0.16-0.19mm) and sealing with clear nail polish.

### Identification and quantification of FRE fungal structures

To identify prevalence of MucFRE-like hyphae, we examined the individual root samples under a compound light microscope at 40 to 100x magnification using pre-established morphological and quantitative parameters (Table 2) (Thippayarugs et al. 1999; Orchard et al. 2017a; Hoysted et al. 2019). Accordingly, we ascribed aseptate hyphae and swellings/vesicles as representative of MucFRE based on the diameter of fungal structures (hyphae<2µm; vesicles<15µm).

**Table 2.** Fine root endophyte (FRE) morphological parameters.

| References | Plant<sup>2</sup> | Hyphal description | Ø (µm) | Vesicle description | Ø (µm) | Arbuscule descriptor |
| --- | --- | --- | --- | --- | --- | --- |
| Rimington et al. 2015<sup>1</sup><br>Hoysted et al. 2019<sup>1</sup> | *Lycopodiella inundata* | Fine | 0.5-1.5 | Terminal/<br>intercalary<br>(pyriform) | 5-15 | Not reported |
|  | *Lycopodiella inundata*<br>(protocorm) | Fine | 0.5-1.5 | Terminal/<br>intercalary<br>(pyriform) | 5-10 | Not reported |
|  | *Holcus lanatus* &<br>*Molinia caerulea* | Smooth fine | 0.5-1.5 | Terminal/<br>intercalary<br>(pyriform) | 5-10 | Arbuscules/<br>arbuscule-like branching |
| Thippayarugs et al. 1999 | *Trifolium subterranean* | Smooth fine | 0.3-0.8 | Not present |  | Hyphal branching |
|  |  | Smooth fine | 0.5-1.5 | Terminal/<br>intercalary<br>(pyriform) | 1.9-3.3 | Hyphal branching |
|  |  | Smooth fine | 0.6-1.4 | Terminal/<br>intercalary<br>(globose) | 3-5 | Hyphal branching |
|  |  | Rough fine | 0.5-1.7 | Terminal/<br>intercalary<br>(pyriform) | 2.6-4.4 | Hyphal branching |
| Orchard et al. 2017a<sup>1</sup> | *Trifolium subterranean* | Fine | < 2.0 | Not reported |  | Fan-like branching |

1- denotes studies with FRE confirmed molecularly as Mucoromycotina
2- roots unless specified otherwise

**Table 3.** Summary of individual roots colonized by site, subplot and plant, comparing spring and autumn 2019.

| Site code | Spring 2019 |  |  |  |  | Autumn 2019 |  |  |  |  |
| --- | --- | --- | --- | --- | --- | --- | --- | --- | --- | --- |
|  | Plants analysed (n) | Roots analysed (n) | Subplots colonised (%) | Roots/plant colonised (%) | Roots colonised (%) | Plants analysed (n) | Roots analysed (n) | Subplots colonised (%) | Roots/plant colonised (%) | Roots colonised (%) |
| TH | 9 | 38 | 67 | 5 | 11 | 7 | 44 | 100 | 70 | 70 |
| HB | 6 | 31 | 75 | 10 | 10 | 11 | 73 | 100 | 73 | 78 |
| ST | 8 | 45 | 75 | 7 | 7 | 11 | 49 | 100 | 78 | 76 |
| PL | 28 | 88 | 57 | 22 | 22 | 19 | 81 | 100 | 85 | 89 |
| CE | 19 | 77 | 100 | 51 | 51 | 19 | 102 | 100 | 99 | 99 |
| MP | 19 | 71 | 0 | 9 | 8 | 17 | 62 | 100 | 98 | 98 |
| BD | 3 | 9 | 50 | 11 | 11 | 16 | 67 | 100 | 98 | 93 |
| MH | 10 | 52 | 80 | 10 | 10 | 8 | 40 | 100 | 75 | 83 |
| SH | 9 | 64 | 20 | 5 | 5 | 12 | 70 | 100 | 80 | 79 |
| NL | 17 | 109 | 0 | 0 | 0 | 23 | 104 | 100 | 87 | 91 |
| AL | 1 | 2 | 0 | 0 | 0 | 3 | 25 | 100 | 56 | 60 |
| **All** | 129 | 586 | 53 | 13 | 14 | 146 | 717 | 100 | 85 | 87 |

In all roots we recorded absence/presence of:

1. MucFRE-like hyphae. Hyphae were identified as present only if observed clearly within the cell walls of at least three root cells. We also noted whether these occurred in root hair cells.
2. MucFRE-like vesicles.
3. Arbuscule-like structures.

‘Coarse’ Glomeromycota-like aseptate hyphae >3µm, if present, were also recorded.

In a subset of colonised roots from two sites representing latitudinal, temperature and precipitation extremes (Coulin Estate (CE), Scotland and Thursley Common (TH), southern England) we measured:

1. Percentage of MucFRE-like hyphal cover. Using an eyepiece micrometer (magnification x63), we subdivided the roots into 250μm sections horizontally and six columns longitudinally (three at each side of the vascular bundle). Percentage cover was calculated as the number of delineated grid boxes containing hyphae (modified from McGonigle et al. 1990 and Sun & Tang 2012) divided by the total number of boxes.
2. Percentage of MucFRE-like hyphal cover attributable to root hair cells, calculated as the number of delineated grid boxes containing colonised hair root cells divided by the total number of colonised boxes.

### Molecular identification

The roots from all 11 sites were processed as above but stored in CTAB lysis buffer (Bainard et al. 2010). We were able to analyse one sample only from Munday Path (MP), Scotland, prior to precautionary closing of the laboratory due to COVID-19.

The DNA extractions were performed according to the method described by Bidartondo et al. (2011) and the fungal 18S region was amplified using the universal primers NS1 (White et al. 1990) and EF3 (Smit et al. 1999). Cloning and sequencing techniques were performed as described in Rimington et al. (2015). Amplicons were cloned (TOPO TA, Invitrogen) and sequenced using an Applied Biosystems Genetic Analyser 3730 (Waltham, MA, USA). Sequences were edited and assembled with Geneious v7.1.9 (http://www.geneious.com) and identified using NCBI BLAST blastn algorithm (Altschul *et al*. 1997).

To expand the number of analysed populations, we added sequences from extracts of *L. inundata* root fungus taken in a previous study (Hoysted et al., 2019) from the same subplots at Thursley Common (TH).

### Analysis of abiotic factors

Monthly temperature and precipitation data for four months immediately preceding sample collection were tabulated for each site. We also analysed these abiotic histories in month 2, 3, and 4 prior to collection to evaluate the temporal evolution of these correlations.

We used 2019 records from the nearest weather stations (www.worldweatheronline.com) as follows: Thursley, Thursley Common (TH); Bere Regis, Hyde Bog (HB); Watergate, Stannon Park (SP); Trenant, Park Lake (PL); Kinlochewe, Coulin Estate (CE); Shieldaig, Munday Estate (MP); Strathcarron, Beinn Damh (BD); Lyndhurst, Matley Heath (MH); Bramshaw, Stadbury Hill (SH); Hampshire, Aldershot (AL); Strabechtse Heide (NL).

### Statistical analysis

#### Measurements of root per plant, root density and root length

To investigate differences in root length, number of roots per plant and root density (root per plant/cm), between sites, a one-way ANOVA was used. Potential correlations between colonization with either root length or root density for each site were tested using the Pearson r correlation test.

#### Quantification of FRE colonization per season and per site

To test the relative contributions of site and season on both the percentage of individual roots, and roots/plant colonised per site, a two-way ANOVA was used, followed by Sidak’s *post hoc* multiple comparisons test. The percentage colonisation was measured as the proportion of total roots evaluated per site, and the weighted average of roots colonised per plant per site. All data was first analysed for normality tests (both Shapiro-Wilk and D’Agostino & Pearson).

Fisher’s exact test was used to compare spring and autumn overall percentages of roots containing FRE vesicle structures. The subsample of roots quantifying colonization within an individual root (percentage of colonization per root) were analysed using unpaired *t*-tests with Welch’s correction.

#### Analysis of abiotic factors

Potential correlations between root colonization per site and local temperature and precipitation histories were tested using Pearson r correlation tests. All statistical tests were analysed using Prism (version 8.4). Statistical significance was established as p value (p) ≤ 0.05.

## Results

Over a six-month growth season, we repeated site visits across all 11 study populations of *L. inundata*. In total we processed and analysed 1,297 roots, 580 from 128 plants in spring and 717 from 146 plants in autumn (Table 2).

### Measurements of root per plant, root density and root length

Mean±SD roots per plant across all sites was 10±2.5 with a density of 2.6±0.7 roots per plant/cm. The mean length of roots was 10.5±2.4mm (Supplemental Table 1). Although there were significant differences between sites for all root measurements (one-way ANOVA, all p < .0001, roots/plant: F_(11,185)_ =3.702; density: F_(11,185)_ =58.21; root length: F_(11,185)_ = 9.89), we found weak correlations between these measurements and fungal colonization (Pearson r, root density: r = −0.146, p = .652; root length: r = 0.238, p = .455).

### Identification of fungal structures

All colonised roots, regardless of season, were predominantly colonised with MucFRE-like hyphae and vesicles/swellings with the exception of Aldershot’s (AL) roots, which had hyphae only but did not have vesicles. Figure 1 shows examples of fine hyphae measuring 0.3-<2µm in diameter with 5-15µm intercalary vesicles and swellings, consistent with the signature hyphal morphotype for Mucoromycotina (Orchard 2017a; Hoysted et al. 2019). Arbuscule-like fine branching structures were present in <0.1% of roots (Figure 1G).

**Figure 1.**
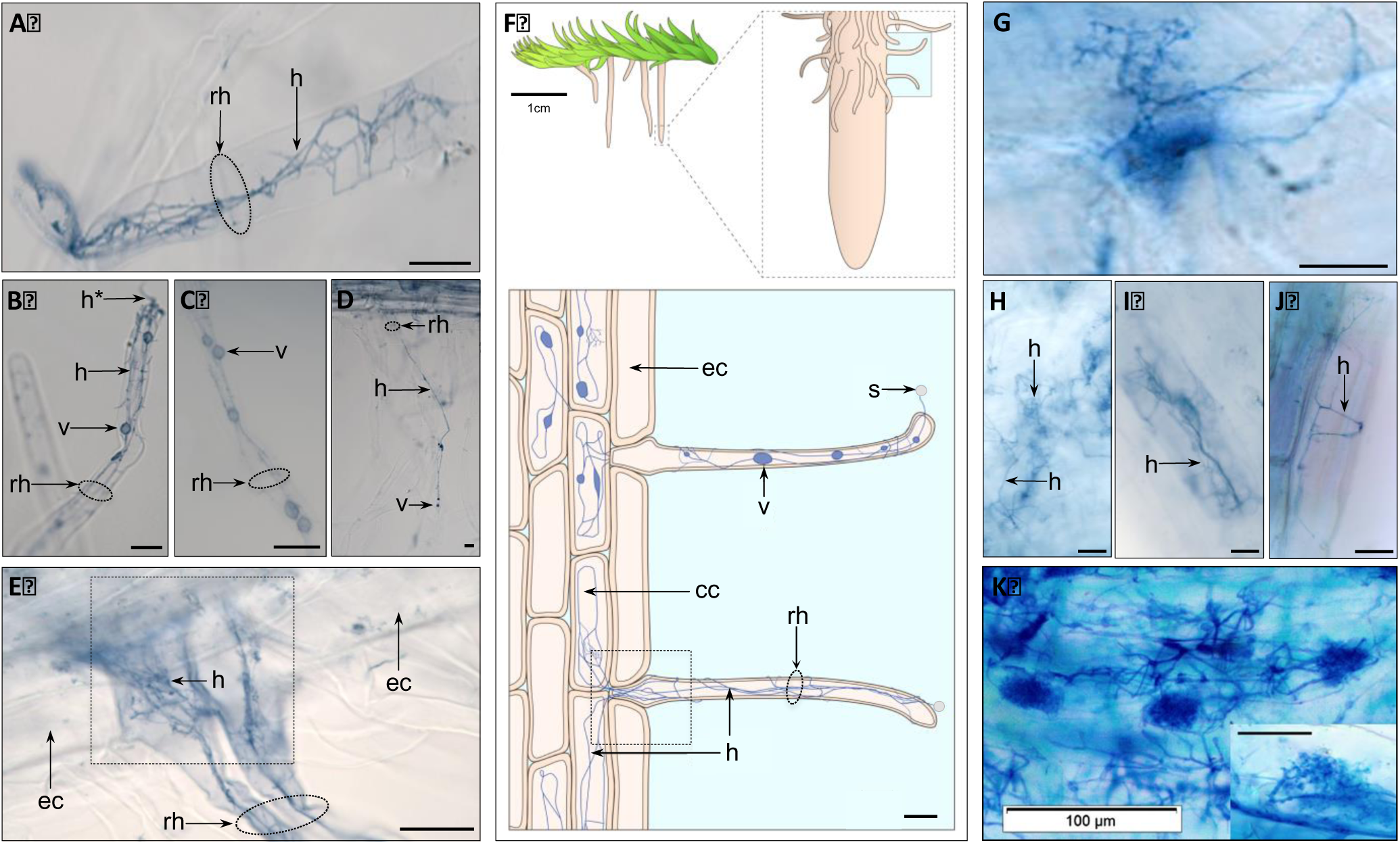
Fine root endophyte (FRE) hyphae with vesicles and arbuscule-like structures in mature *Lycopodiella inundata* sporophytes. Both root hair cells (A-E) and roots (G-J) showing examples of fine hyphae and intercalary vesicles or hyphal swellings ranging 2-10µm. Some hyphae were seen entering through the root hair tip (labelled h* in B). (E) Two adjacent root hair cells with bundles of FRE strands twisting and branching throughout and colonizing cortical cells but skipping epidermal cells. (F) Schematic sketch of a *Lycopodiella inundata* plant illustrating a root, root hair cells, fine hyphae, vesicle, spores and epidermal and cortical cells (expanded shaded box on bottom). The dotted square in (E) highlights a root hair position between epidermal cells renderized in the equivalent dotted box of the sketch. Arbuscule-like branching were also seen (G). FRE were observed branching and twisting throughout the root (H), individual epidermal root hair cells (I) and cortical cells (J). (K) Previously published FRE and arbuscules (inset) in *Trifolium subterraneum* root (adapted from Orchard et al. 2017a *New Phytologist* with permission). All micrographs are acidified Schaeffer blue ink. Labels: ‘rh’ root hair; ‘h’ fine hyphae; ‘v’ intercalary vesicle; ‘s’ spore; ‘cc’ cortical cell; ‘ec’ epidermal cell. All scale bars 20µm, unless detailed otherwise.

Between 2% and 10% of the roots from Dutch and southern English sites had wider knobbly aseptate hyphae appearing AMF-like (Figure 2). Of these, only a minority (1%) of the roots had AMF-like vesicle structures (Figure 2C,D,G). We did not observe this hyphal morphotype in Scottish roots.

**Figure 2.**
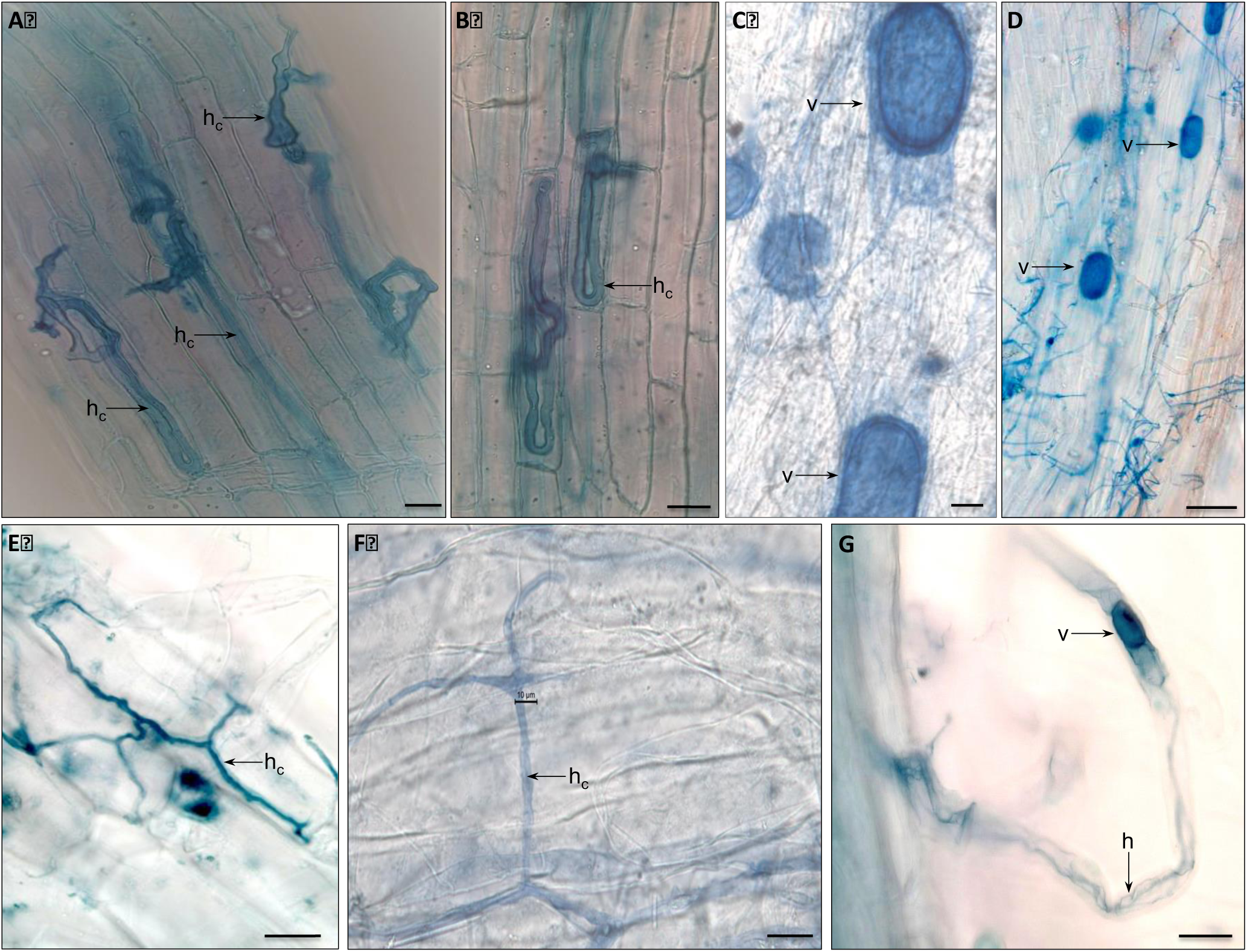
Arbuscular mycorrhizal-like vesicles and ‘coarse’ aseptate hyphae. (A-C,E,F) Acidified Schaeffer blue ink light micrographs of mature *Lycopodiella inundata* sporophyte root showing intracellular (A,B) and intercellular (C,E,F) coarse hyphae ‘h_c_’, as seen in 4% of colonised roots in autumn. (C) Large vesicles ‘v’ up to 80µm. (D) Reference micrograph of *Holcus lanatus* root colonised by both Mucoromycotina fine root endophyte hyphae ‘h’ and Glomeromycotina v; adapted from Hoysted et al. 2019 (Copyright American Society of Plant Biologists). (G) Detail of colonised root hair with fine hyphae and large Glomeromycotina-like vesicle. Labels: ‘h’ fine hyphae; ‘h_c_’ coarse hyphae; ‘v’ vesicle. All scale bars 20µm.

### Quantification of fungal structures

#### Presence of MucFRE-like hyphae

We found statistically significant differences in MucFRE-like hyphal presence in spring vs. autumn in both season and site factor in the percentage of colonised individual roots/site (two-way ANOVA, season: F_(1,1250)_ = 1148, p < .0001; site: F_(9,1250)_ = 15.85, p < .0001). The interaction term was also significant indicating the effects of season and site are dependant on each other (two-way ANOVA, season: F_(9, 1250)_ = 6.238)(Figure 3, Table 2).

**Figure 3.**
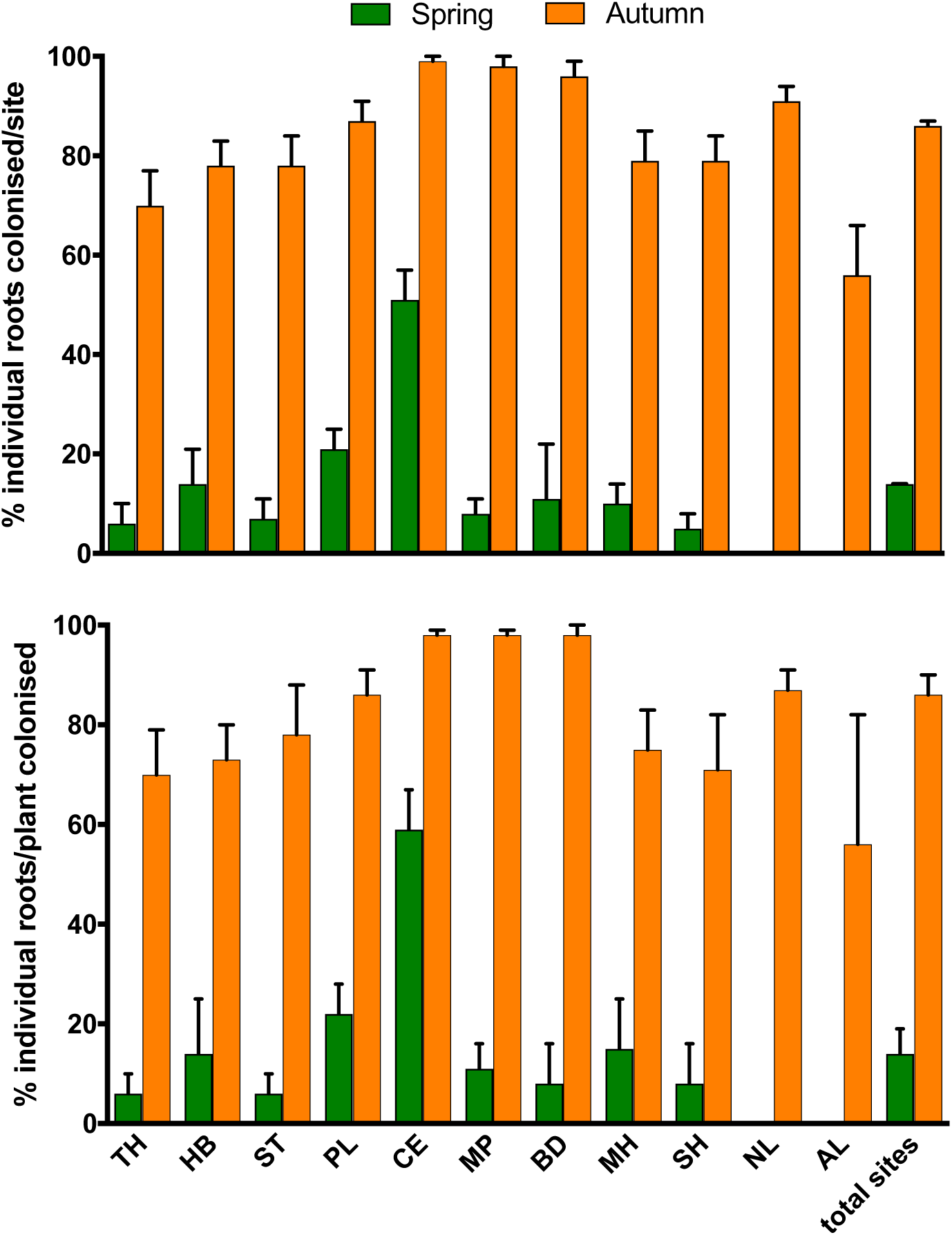
Comparison of spring and autumn 2019 roots colonised by fine root endophytes across all sites (except Aldershot). Top) percentage of colonised individual roots by site (two-way ANOVA, season: F_(1,1250)_ = 1148, p < .0001; site: F_(9,1250)_ = 15.85, p < .0001). Bottom) percentage of colonised individual roots per plant by site (two-way ANOVA, season: F_(1, 251)_ = 363.6, p < .0001; site: F _(9, 251)_ = 6.788, p < .0001). Values for n for each site are shown in Table 2. Error bars ±SE. There are no bars in spring for both AL_(n=2 roots)_ and NL_(n=109 roots)_ as they had no colonised roots.

We also found significance in both the season and site factors when looking at percentage of roots colonised/plant (two-way ANOVA, season: F_(1, 251)_ = 363.6, p < .0001; site: F_(9, 251)_ = 6.788, p < .0001). As above, the interaction term was also significant (two-way ANOVA, season: F_(9,251)_ = 2.8, p = .004) suggesting regarding percentage of roots per plant, the season effect was not the same across all sites. Scotland’s Coulin Estate had significantly more colonization per site (51%) in spring compared to all other sites, including the other two geographically close Beinn Damh (11%) and Munday Path (8%) sites. Note that AL was excluded from this analysis due to the low number of roots sampled (n = 2 in spring).

Overall 14% of the individual roots were colonised in spring vs. 86% in autumn. Regardless of season, all colonised roots had FRE present in root hair cells.

#### Presence MucFRE-like vesicles and swellings

Vesicles and swellings were significantly more prevalent in the autumn vs. spring for all sites. Overall, vesicles were present in 8.8% of total roots analysed in the autumn, as opposed to 2.4% in the spring (Fisher’s exact test, p < 0.0001). Arbuscule-like structures were observed in <1% of the autumn-collected roots only (Figure 3G).

#### MucFRE-like hyphal colonization

MucFRE-like hyphal colonisation was analyzed in a subset of n=32 colonised roots from CE and TH. The percentage of an individual root colonised by FRE was significantly different in spring and autumn for CE (Scotland) but not TH (southern England), however only two TH roots were colonised in spring. Mean±SD root FRE area coverage for CE was 10.3±1.7% in spring and to 33.5±15.79 in autumn (Welch’s *t*-test, t = 4.295, df = 11.23, p = .001). Mean±SD root FRE cover for TH was 35±21.2% in spring and to 22.23±18.31% in autumn (Welch’s *t*-test, t = 0.199, df = 3.402, p = 0.853).

Hyphae typically occurred above the root cap and showed a marked propensity to occur in root hairs. Root hair colonization contributed to 43.7% of the total root colonization in autumn and 28.8% in spring.

### Molecular identification

The OTUs analysed from a representative sample from Munday Estate (MP) in Scotland was consistent with Mucoromycotina species complex and aligned best with isolate BVMT_30 (sequence ID: MH174565.1). Three Mucoromycotina fungi OTUs were also detected by Hoysted et al. (2019) in wild *L. inundata* population roots collected from the same TH subplots sampled in this study.

### Abiotic factors

Temperature and precipitation data for 2019 showed significant differences across sites. The Scottish sites had the highest rainfall both annually and monthly, and the Dutch site and Hyde Bog the lowest. We also noted peak rainfall occurring two months earlier for Scotland than all the other sites. The lowest monthly and annual average temperatures occurred in Scotland.

The correlation graph (Fig. 4) shows mean temperature for the 30d preceding root sampling had a strong negative correlation with percentage of roots colonised at sampling for both seasons (spring: r = −0.81, p = .003; autumn: r = −80, p = .003). For the cumulative precipitation 30d preceding root sampling, there was a strong positive correlation with the autumn roots but only a week tendency with the spring roots (spring: r = 0.32, p = .339; autumn: r = 0.74, p = .008). We found the correlation strength diminishing in relation to the time series data. By four months preceding sampling, correlations were not significant (Figure 5).

**Figure 4.**
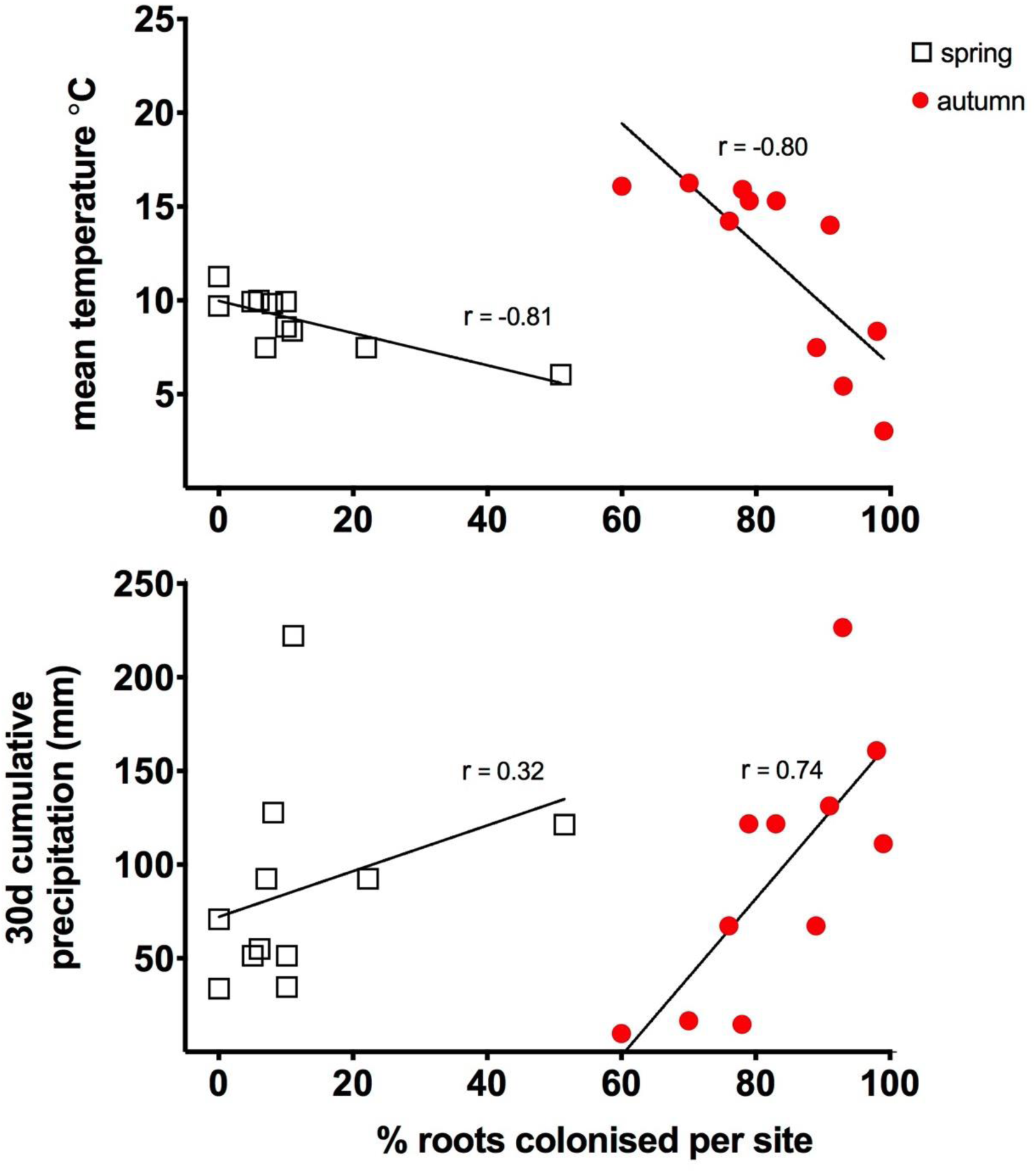
Correlation between abiotic factors and colonised roots per site per season. Top) mean temperature 30d preceding root sampling (Pearson r, spring: r = −0.81, p = .003; autumn: r = −80, p = .003). Bottom) cumulative precipitation 30d preceding root sampling (spring: r = 0.32, p = .339; autumn: r = 0.74, p = .008).

**Figure 5.**
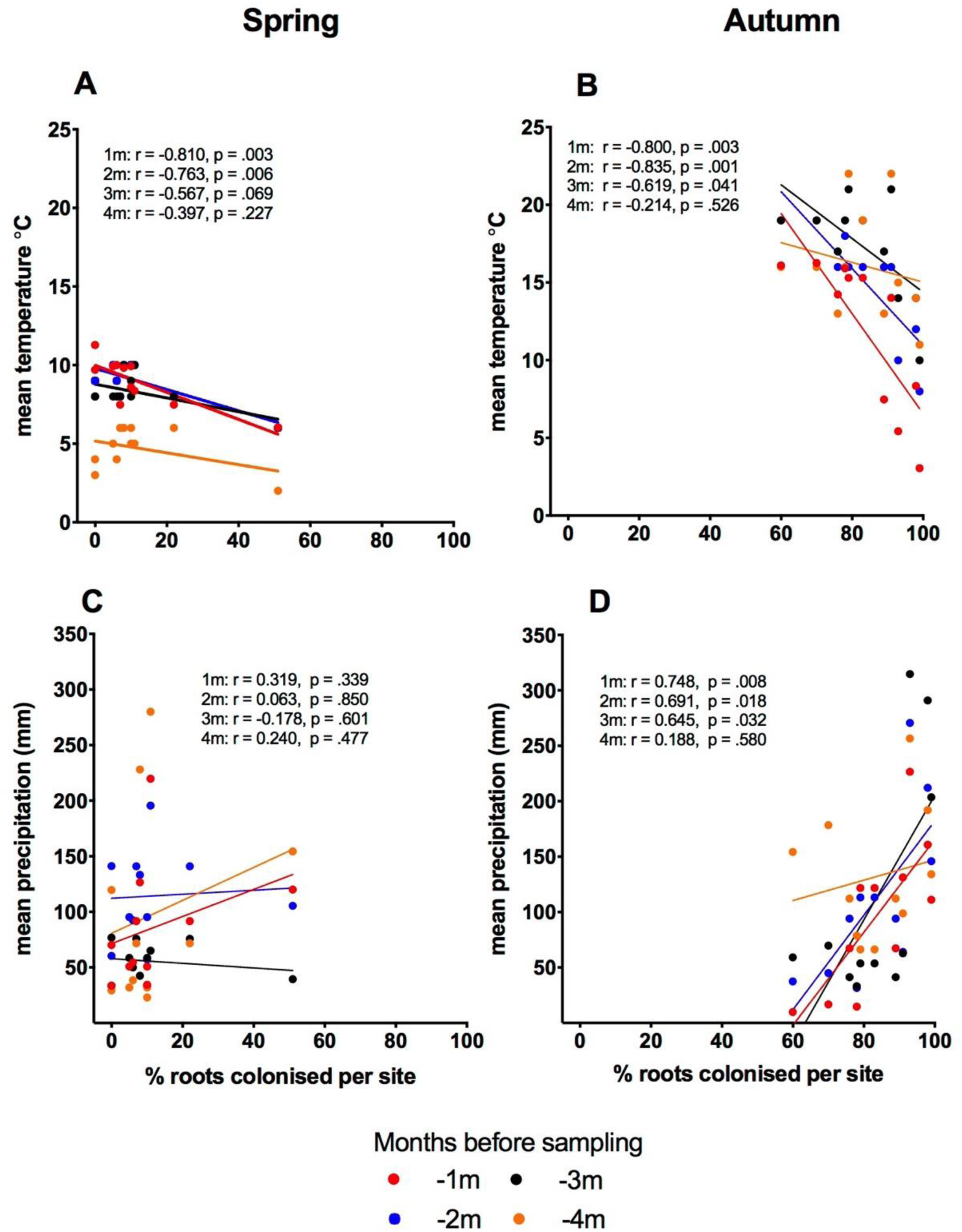
Correlation time series between abiotic factors and individual colonised roots per site for the four consecutive months preceding sampling (spring and autumn). Root colonization correlated strongly with temperature in spring and autumn (A/B) and precipitation (C/D) in autumn only. Observed correlations were stronger at 0-30d or 31-60d and progressively weakened thereafter. ‘m’ = months.

## Discussion

### Plant phenology is an indicator for fungal colonization

We detected significant differences in the percentage of FRE colonization in individual roots/site and colonized roots/plant at the beginning of the host plant’s growing season compared with the end. These differences were seen across all 11 sites. Some sites presented more dramatic differences than others, e.g. in the Netherlands no roots were colonised by MucFRE-like hyphae in the spring but 87% were colonised in the autumn. This seasonal change most likely relates to climate history because there were correlations between root colonization and site-specific temperature and precipitation histories in the months leading up to sampling collection.

Our study confirms plant phenology has a strong influence on fungal colonization. In light of our results, we would suggest caution in the interpretation of previous ecological studies examining root colonization from samples collected at a single time point (Urcelay et al. 2011; Bueno de Mesquita 2018b; Pereira et al. 2020). Conclusions on the potential influence of edaphic or environmental variables in colonization detected may have been related more to seasonality than to a true influence of these variables. Sampling and analysis following hyphal dormancy (Kabir et al. 1997) may also explain why some plants had previously been classified as ‘non-mycorrhizal’, e.g. *Buddleja davidii* (Dickie et al. 2007) or yielded a low overall mycorrhizal rate based on fungi identified molecularly, e.g. 13% of previously sampled *L. inundata* roots (Rimington *et al*. 2014).

There are numerous reports studying phenology of AMF, particularly in grasslands (Bohrer, Amon and Friese 2004; Escudero and Mendoza 2005; Lingfei 2005; Mandyam and Jumpponen 2008) and agricultural fields (Kabir et al. 1997; Saif and Kahn 1975; Tian 2011), in contrast with the scarcity of studies examining FRE phenology. A recent meta-analysis of temporal changes in AMF and FRE colonization (Bueno de Mesquita et al. 2018a) found that 75% of the studies detected temporal changes over the growth season. However, the inclusion of FRE as distinguished taxonomically from AMF was not clear.

In the single previous mycorrhizal phenology study including *L. inundata*, more colonization was found in the spring than autumn (Fuchs and Haselwandter 2004), in sharp contradiction to our results. The explanation for this different result is not clear to us but it may relate to a small sample group fortuitously biasing the results, lack of separation between FRE from AMF during analysis, and/or lack of molecular identification of fungi.

In a single-site experiment examining fungal colonization of four forb species (*Polygonum bistortoides, Gentiana algida, Artemisia scopulorum* and *Geum rossii)* over a three-month alpine growing season, Bueno de Mesquita et al. (2018a) demonstrated colonization (potentially of FRE, as well as AMF and dark septate endophytes) peaked as the angiosperm fruiting began and AMF vesicles increased as plants produced seeds. Soil temperature and moisture, and plant phenology contributed to root colonization levels, depending on plant species. They also found fungal propagules from the soil colonized new roots within days. The FRE colonization, including vesicles in *L. inundata*, may also be peaking at the onset of strobilus formation. In fact, all 11 populations had produced strobili by September/October before we collected our samples. In another FRE colonization study using pot cultures, significant differences in *Trifolium subterraneum* FRE colonization were documented at the beginning and end of the growing season (Thippayarugs 1999).

### Colonization differences correlated with temperature and precipitation

Colonization differences across study sites correlated with temperature and precipitation data suggesting local environmental variables are contributing to phenology differences. Interestingly, the two sites with significantly higher presence of hyphae per site in the spring, Coulin Estate (CE), compared with all other sites also had the lowest monthly temperatures in the 30d leading up to sampling; CE (mean = 6.2**°**C), suggesting FRE may have preferential temperature regimes triggering growth. The sites with temperature extremes leading up to the autumn collections, Aldershot (AL) with amongst the highest (mean = 16.2**°**C) and CE with the lowest (mean = 3.1**°**C), had an inverse relationship with colonization; temperatures 30d and two months leading up to sampling was more strongly correlated than at three or four months. Our correlation time series analysis further supports the possibility that hyphal growth may be initiated or inhibited at certain temperature limits. Notwithstanding, whether this correlation also implies causation cannot be ascertained from this field study.

Precipitation histories for spring did not correlate to FRE root colonization, however the months leading up to the autumn collection did correlate. This might suggest precipitation is more important to FRE activity when temperatures are hotter.

Therefore, one can only speculate whether AL has the most vulnerable population across the study may be related to its higher temperatures and lower precipitation rates than the other sites.

### Other biotic and abiotic factors contributing to colonization

There are several other factors that may influence FRE root colonization including atmospheric N deposition, plant community structure, soil chemistry composition or other unseen biotic and abiotic dynamics at the local level. Nitrogen loads are known to be much higher in the Netherlands and much of southern England (de Heer et al. 2017, Lilleskov et al. 2019) compared to northern Scotland and can have an effect on mycorrhizal fungi (Ceulemans et al. 2019). None of the spring root samples from the Netherlands were colonised compared with 51% at CE Scotland. This extreme may also be attributed to precipitation and temperature histories, but local differences in N deposition cannot be ruled out. The other two nearby Scottish sites had significantly lower spring colonization rates of 9% (MP) and 11% (BD), more in line with the other sites, suggesting local edaphic conditions may have also played a role in CE’s higher colonization rates. Only field-based experiments testing wet and dry N with various N loads will be able to provide further insight into FRE responses over the growth season as well as the impact on FRE host-plants.

Studying these conditions more closely could help determine which environmental conditions are better suited for both plant-host and FRE guiding conservation planning. This is a critical point with respect to *L. inundata* conservation efforts in warming regions of Britain, particularly with lower precipitation. Until we understand these factors better, we suggest population restoration and conservation efforts focus on areas of the country with more suitable temperature and precipitation regimes.

Vegetation community composition could play a role in belowground microbial competition (van der Heijden et al. 1998, Wardle 2004). Other biotic and abiotic dynamics could also be relevant. For example, plant strigolactones exudation plays a role in triggering Glomeromycota hyphal branching *in vitro* (Akiyama et al. 2005; Besserer et al. 2006). However, their ecological role for plant-host fungal preference has not been definitely determined (Mishra et al. 2017; Carvalhais et al. 2019). Although we did not compare root colonization cover per root among all 11 sites, if FRE signalling and development resemble those of AMF, we expect the extent of reciprocal nutrient exchange (Smith and Smith 2011), e.g. if correlated with the percentage of the roots colonised at any one time, to be governed locally by host-plant P (Karandashov and Bucher 2005, *Smith et al. 2015*) and N requirements (Hoysted et al. 2019). Conversely, the plant may have an excess of photosynthate resources, which it can opportunistically provide to compatible symbionts. This will vary given environmental conditions, competition for surplus resources and ‘sink strength’ (Walder and van der Heijden 2015) and we expect requirements will certainly shift over the host plant’s lifespan (Field et al. 2015b). The fungal carbohydrate and lipid requirements (Jiang et al. 2017; Luginbuehl et al. 2017) of MucFRE may also contribute to colonization responses. Further studies will be necessary to tease apart the contribution of the variables affecting FRE colonization of *L. inundata* and host plant retention.

### Lycopodiella inundata’s preferential association with FRE

The overwhelming majority of FRE observed in mature sporophyte roots exhibited typical Muc-FRE traits, suggesting *L. inundata* may have a rarely seen plant preference for an endomycorrhizal fungus (Walder and van der Heijden 2015). Interestingly, we also discovered that every colonised root had FRE present in the root hair cells. Whether the thinner root hair cell walls indicate a preferential entry point zone for FRE will require electron microscopic analyses.

The low number of observed MucFRE-like arbuscules in *L. inundata* in our study is consistent with previous reports (Hoysted et al. 2019). Given the ephemeral nature of arbuscule development (Roth et al. 2019), one possibility is that they had already developed and senesced before strobilus production, and had decomposed before we collected roots at the end of the growing season.

Although we obtained molecular confirmation in one site, and roots from the same subplots on Thursley had previously been identified as MucFRE, we cannot completely confirm the rest are also MucFRE without further molecular analysis across all remaining sites. However, the possibility of finding different taxa with the same anatomical features as MucFRE is highly unlikely.

We also noted a minority of the sporophyte roots contained aseptate hyphae with diameters up to 2.5µm (4% were >3µm). These AMF-like hyphae were seen in a small fraction of roots in the lowland heathland sites, but not in Scotland. This may represent multiple FRE species interacting (Thippayarugs 1999; Orchard 2017a) or rare AMF co-colonization, either opportunistic or driven by plant nutrient requirements.

## Conclusion

In this large-scale and intensive study, MucFRE-like hyphae were overwhelmingly present at the end of the season - colonizing 86% of roots compared with 14% in the spring - confirming a strong phenological pattern for mycorrhizal fungi, at least in *L. inundata*. Appreciating MucFRE presence does not directly convey functionality (Cosme et al. 2018), previous studies incorporating symbiotic functional responses may have underestimated potential nutrient exchange between fungus and plant due to harvest time of source host-plants. This may also be pertinent to agricultural studies measuring colonization and growth responses to different treatments.

Although seasonal variation in mycorrhizal root colonization has not been observed in perennial woodland plants (Brundrett and Kendrick 1988), the fact that collection of samples for large-scale ecological studies often occurs over several months also warrants caution in interpreting results as season may play a strong role. For plant species habitat conservation plans, the success of a particular plant-fungus mutualism can make or break survivability. Our results strongly indicate that studies of mycorrhizal fungal species composition or colonization rates must be designed and evaluated taking phenology as a crucial variable.

## Acknowledgements

We gratefully acknowledge financial and institutional support from The Royal Botanic Gardens, Kew. Valuable time and logistical support were provided by the private and quasi-public land owners/managers including Staatsbosbeheer, Brabant-Oost, Natural England, the New Forest, National Trust and Southwest Water, all of whom welcomed the research and assisted with site access and/or permits. Special thanks to Drs. Jeffrey Duckett, Silvia Pressel and Katie Field who provided valuable comments during the manuscript development, Dominic Price, Species Recovery Trust, for helping identify viable sites and Dr. Susan Jarvis, UK Centre for Ecology and Hydrology, who reviewed the statistical tests and design of the study. Additionally, the many students who helped process the roots in the laboratory and especially Victoria Jaggers who contributed the drawing for Figure 2.

## Consent for publication

Permissions to reprint two Figure images were granted and relevant copyright is stated in the figure legend.

## Authors’ contributions

JK conceived and designed the study, refined the root staining and colonization assessment protocols, examined the roots for colonization, compiled and analysed the data, and wrote the manuscript. All co-authors contributed to the manuscript. More specifically, EA stained and examined roots and conducted molecular analyses to identify the fungus, MB supervised the molecular analyses and JS assisted all fieldwork and data analysis.

